# AlphaGenome-enabled analysis of non-coding regulatory variants underlying *RHD* expression with wet-lab validation

**DOI:** 10.64898/2026.01.21.700828

**Authors:** Miao Liu, Ziyang Shen, You Kyeong Jeong, Na Yu, Shang-Chuen Wu, Andre Wittig, Danielle Tenen, Yanjing Liu, Jun Liu, Li Chai

## Abstract

Systematic identification of functional non-coding regulatory variants remains a major challenge in human genetics. Conventional approaches such as large-scale CRISPR screening and genome-wide association studies (GWAS) are powerful but often prohibitively expensive, time-consuming, and experimentally intensive, limiting their scalability for locus-specific mechanistic studies. Recent advances in artificial intelligence offer the potential to partially replace or substantially augment these approaches by prioritizing regulatory variants with high functional likelihood. The *RHD* antigen, a major contributor to red blood cell alloimmunization, hemolytic transfusion reactions, and hemolytic disease of the fetus and newborn, serves as an excellent model for this paradigm, since coding variants alone do not fully account for differences in *RHD* expression.

Here, we present an integrated artificial intelligence (AI)-guided and experimental framework to identify and validate functional non-coding regulatory variants governing *RHD* expression. We first applied AlphaGenome (AG), a deep-learning model released in 2025 for non-coding variant impact prediction, to systematically interrogate the *RHD* locus. By integrating multi-omics datasets, AG prioritized regulatory regions within the promoter, 5′ untranslated region (5′UTR), and intragenic regions. In silico deletion- and Single Nucleotide Polymorphism (SNP)-based perturbation analyses consistently predicted that variants within the promoter and its proximal regions, as well as within intragenic regions, exert strong suppressive effects on *RHD* expression.

To experimentally validate these predictions, we performed CRISPR-mediated base editing in K562 cells at AG-prioritized non-coding SNP sites. Editing of a high-score predicted variant (chr1:25272434 G>A) achieved efficient base conversion and was accompanied by additional nearby edits, all predicted by AG to downregulate *RHD* expression. In contrast, editing of low-score predicted sites (chr1:25272422 C>T) produced much smaller functional effects. Quantitative polymerase chain reaction (qPCR) analysis of full-length *RHD* transcripts, together with flow cytometry–based analysis of *RHD* expression, confirmed strong concordance between AI-based predictions and transcriptional as well as phenotypic outcomes.

Taken together, our results demonstrate that the combination of AI-guided regulatory variant prioritization and targeted base editing provides a potentially scalable and cost-effective alternative to traditional CRISPR screening for decoding functional non-coding variants in blood group genes, with direct implications for genomics-based RHD typing and transfusion medicine. To our knowledge, this study also represents the first validation of AlphaGenome predictions at the phenotypic level using wet-lab experiments.

## Introduction

The functional interpretation of non-coding genetic variation remains one of the most significant unresolved challenges in modern genomics. Although large-scale approaches such as genome-wide association studies (GWAS) and CRISPR-based screening have transformed our understanding of gene regulation and disease susceptibility, these methods are often constrained by high cost, extensive experimental burden, and limited resolution for locus-specific regulatory mechanisms. ^1,2^ Genome-wide CRISPR screens typically require the construction and validation of large guide RNA libraries, specialized cell models, and deep sequencing infrastructure, while GWAS demands massive sample sizes to achieve sufficient statistical power and frequently yields associations that are difficult to mechanistically interpret. As a result, there is a growing need for alternative strategies that can efficiently prioritize functional regulatory variants while reducing experimental complexity and cost.^1,3^

Recent advances in artificial intelligence (AI) and deep learning have introduced new opportunities to address this gap. By integrating diverse epigenomic, transcriptomic, and chromatin conformation datasets, AI-based models have begun to predict the functional consequences of non-coding variants on gene expression with increasing accuracy. Rather than replacing experimental approaches entirely, these models offer the potential to guide and refine downstream validation, enabling focused interrogation of high-probability regulatory elements and substantially reducing the scale of wet-lab screening.^4,5^ Such an AI-assisted paradigm is particularly well-suited for genes with complex regulatory architectures and strong clinical relevance, where traditional approaches have struggled to fully explain phenotypic variability.

The *RHD* blood group antigen represents a compelling example of this challenge. Encoded by the *RHD* gene on chromosome 1, *RHD* is one of the most clinically significant erythrocyte antigens, second only to ABO in transfusion medicine. The presence or absence of *RHD* expression determines Rh-positive or Rh-negative status and plays a central role in hemolytic transfusion reactions, hemolytic disease of the fetus and newborn, and alloimmunization.^6,7^ Even subtle quantitative differences in *RHD* expression can have important clinical consequences, influencing serologic typing accuracy and transfusion compatibility.^8^ Importantly, transfusion-related complications contribute substantially to morbidity and clinical outcomes across a broad spectrum of diseases.^9,10^ Accurate genotyping of *RHD*, together with other clinically relevant red cell antigens, is therefore becoming increasingly important as red blood cell (RBC) genotyping is more widely adopted in transfusion medicine.^,11^ Genotype-based donor–recipient matching represents an emerging direction for improving transfusion safety and efficacy, particularly in patients requiring repeated transfusions. Several European countries, including Germany and the Netherlands, and pilot initiatives in the United Kingdom, have already begun exploring the use of next-generation sequencing (NGS) for large-scale donor typing.^11-13^

Over the past several decades, extensive effort has been devoted to characterizing coding variants within *RHD*. Missense mutations, nonsense mutations, splice-site alterations, and gene rearrangements have been linked to weak D, partial D, and DEL phenotypes, and these coding alterations are now routinely incorporated into molecular blood group typing strategies. However, coding variants alone fail to account for a substantial proportion of observed variability in *RHD* expression. For example, while nearly all *RHD*-negative cases in Caucasian populations can be explained by complete *RHD* deletion, this paradigm does not hold across other ethnic groups. In a large cohort of 2,493 serologically D-negative donors from China, 67.6% carried *RHD* deletions and 7.4% harbored nonfunctional *RHD* alleles caused by exonic frameshift or hybrid events, indicating that approximately 75% of cases could be attributed to clear coding-region loss-of-function. Notably, up to 25% of *RHD*-negative phenotypes in this population were associated with non-coding alterations, predominantly affecting splicing, underscoring the substantial contribution of regulatory variation to *RHD* expression heterogeneity. Individuals with identical coding sequences can display markedly different serologic phenotypes, suggesting that regulatory mechanisms beyond the coding region play a critical role in determining *RHD* transcription and protein abundance.^14^

Non-coding regulatory regions—including promoters, untranslated regions (UTRs), introns, and distal regulatory elements—are increasingly recognized as key determinants of gene expression. In other biological systems, variants within these regions have been shown to alter transcription factor binding, chromatin accessibility, RNA processing, and transcript stability, leading to clinically meaningful phenotypes.^7,15^

Several factors have contributed to this gap. First, the *RHD* locus resides within a complex genomic region characterized by high sequence homology with RHCE, making variant calling and functional dissection technically challenging.^6^ Second, erythroid-specific regulation adds another layer of complexity, as regulatory elements may be active only in defined stages of erythroid differentiation.^16^ Finally, experimental validation of non-coding variants traditionally requires labor-intensive reporter assays or large-scale CRISPR perturbation screens, approaches that are difficult to deploy comprehensively across extensive regulatory regions.^17-19^

AI-based regulatory variant prediction models offer a promising solution to these challenges. By learning patterns from large-scale functional genomics datasets—including RNA sequencing, chromatin accessibility assays, histone modification profiles, and three-dimensional genome organization—such models can infer how sequence changes are likely to influence gene expression in specific cellular contexts. Importantly, these predictions can be generated at single-nucleotide resolution, enabling fine-grained prioritization of candidate variants for experimental follow-up.^20,21^ For genes like *RHD*, where the regulatory architecture is complex, but the clinical stakes are high, AI-guided prioritization provides an opportunity to move beyond descriptive genomics toward mechanistic understanding. Against this backdrop, AlphaGenome (AG) provides a new technological foundation for the systematic interrogation of non-coding regulatory variants. AG is a deep learning–based genome function prediction model released in 2025, designed to infer the potential impact of non-coding sequence variation on gene expression and regulatory states directly from DNA sequence. The model is trained on large-scale, integrated multi-omics datasets, including RNA sequencing, chromatin accessibility (Assay for Transposase-Accessible Chromatin using sequencing, ATAC-seq), histone modification profiles, transcription factor binding, and three-dimensional chromatin organization. This architecture enables single-nucleotide–resolution prediction of regulatory consequences arising from sequence perturbations. Unlike conventional approaches that rely on isolated epigenomic features, AG adopts a unified modeling framework that jointly captures multiple layers of gene regulation, thereby improving predictive performance in complex regulatory regions such as introns and promoter-proximal elements. Importantly, AG supports expression-effect prediction across multiple human cell types and tissue contexts, encompassing dozens of cell lines and differentiation states.^22^ This capability allows regulatory variants to be evaluated in a cell-type–specific manner, which is particularly critical for genes with highly context-dependent regulation, such as the erythroid-specific expression of *RHD*. Collectively, these features enable a direct and scalable connection between AI-based prioritization of non-coding variants and targeted experimental validation.

In this context, erythroid cell models provide a tractable system for linking regulatory variation to functional outcomes. Cell lines such as K562 and HUDEP2 recapitulate key aspects of erythroid gene regulation and allow direct measurement of *RHD* transcripts and surface antigen expression. When combined with precise genome editing technologies, including base editing, these systems enable causal testing of individual non-coding variants without introducing large-scale genomic disruptions.^23,24^

These considerations highlight a critical unmet need in transfusion medicine: a scalable, mechanistically informed framework to identify functional non-coding regulatory variants that govern *RHD* expression. Bridging AI-based prediction with targeted experimental validation offers a path forward, complementing existing GWAS and CRISPR-based approaches while mitigating their limitations. A deeper understanding of *RHD* regulatory architecture will not only improve molecular blood group typing but also provide a generalizable blueprint for studying non-coding variation in other clinically important blood group systems.

## Methods

### Definition of the AlphaGenome Screening Window for *RHD* regulation

To define the AG screen window, *RHD* regulatory regions were manually predicted by integrating histone marks, chromatin accessibility, CpG features, enhancer annotations, and in-house DNA methylation data on the UCSC Genome Browser (GRCh38/hg38). Public ENCODE-derived tracks (H3K4me3, H3K4me1, H3K27ac, DNase I hypersensitivity, CpG islands, GeneHancer) from leukemia cell line K562 (with *RHD* expression) cells were overlaid with custom liver cancer cell line SNU398 (without RHD expression) methylation and CpG-density tracks, and all candidate element coordinates were recorded in hg38 and cross-checked against RefSeq/GENCODE *RHD* exon–intron annotations to ensure accurate mapping of promoter and intragenic regions. Unless otherwise specified, genomic coordinates were reported at the *RHD* gene locus level (hg38). For base editing experiments, target sites were designed based on the *RHD*-201 transcript structure.

For AG in silico prediction, the primary screening window for *RHD* was defined as a genomic interval extending from 10 kb upstream of the annotated transcription start site (TSS) to 10 kb downstream of the 3′ end of the *RHD*-201 transcript. This design was chosen to (i) capture the full promoter and promoter-proximal region upstream of the TSS, where clustered H3K27ac, H3K4me3, H3K4me1, and DNase I hypersensitivity peaks in K562 cells indicated a compact zone of *RHD*-focused regulatory activity, and (ii) include intragenic and immediately downstream non-coding segments, which showed persistent H3K27ac/H3K4me1 signal and are increasingly recognized as sites of intronic or 3′-proximal enhancers and silencers. Although additional regulatory marks were observed at greater distances, these more distal peaks frequently overlapped domains assigned to neighboring genes (such as RHCE), lowering the prior probability that they represent *RHD*-specific elements and motivating their designation for secondary analyses rather than primary AG interrogation.

### In silico prediction of non-coding variant effects using AlphaGenome

AG (research preview version released by Google DeepMind in July 2025) was executed in a Google Colab environment using Python scripts through the official AG research notebook interface. ^22^ The human reference genome hg38 was used for all analyses. Although AG supports the prediction of variant effects across multiple cellular contexts, all analyses in this study were performed in the K562 cell, which expresses *RHD*. RNA-seq was used as the output readout to precisely capture the relationship between genetic variation and *RHD* gene regulation.

For genome interval prediction, AG retrieves the reference genome from cloud storage and extracts all annotated transcripts using the built-in transcript_extractor. Transcripts associated with *RHD* were subsequently selected to define the detailed genomic architecture of the gene. However, AG supports predictions only for specific DNA sequence lengths; therefore, all analyzed genomic intervals were constrained to supported sizes (2 kb, 16 kb, 100 kb, 500 kb, or 1 Mb), ensuring that the *RHD* locus was fully contained within the selected sequence window.

For variant effect prediction, the score_variant function was used to evaluate the impact of genetic variants on *RHD* expression. For RNA-seq outputs, AG computes a variant effect score (raw score) based on the predicted expression difference (log-fold change) between reference (REF) and alternative (ALT) sequences. Scores below zero were interpreted as downregulation of *RHD* expression, with the absolute value of the score reflecting the magnitude of the effect. Candidate variants were ranked according to their predicted scores, and the top ten variants were selected independently from intronic regions and from regions spanning ±10 kb around the TSS for downstream experimental validation (Figure 2).

The analyses included: (i) definition of the *RHD* predictive regions and baseline expression scores; (ii) perturbation analyses of intragenic non-coding regions sequences through base deletions or SNPs; (iii) SNP and deletion analyses within the ±10 kb TSS-flanking regions; and (iv) design of base-editing target sites, together with quantitative PCR and flow cytometry–based readouts to evaluate *RHD* expression.

### Selection algorithm for variant validation

To contextualize AG-predicted non-coding variants within existing *RHD* genetics knowledge base, all candidate sites were systematically cross-checked against published literature and reputable Rh databases. First, each AG-prioritized variant was queried in PubMed and major transfusion medicine journals to identify prior case reports or functional studies describing its association with *RHD* phenotypes. In parallel, variants were compared against entries in RhesusBase and the International Society of Blood Transfusion (ISBT) Rh allele tables to determine whether they mapped to known *RHD* alleles or previously cataloged regulatory changes.^25^ Finally, population allele frequencies were retrieved from large-scale reference datasets (e.g., GNOMAD) to assess the presence and frequency of each variant across diverse ancestries. Variants selected for initial experimental validation were required to meet three criteria: (i) a high predicted downregulatory effect on RHD expression by AG; (ii) presence in at least one population database, preferably at low allele frequency consistent with a rare or underrecognized regulatory allele; and (iii) absence of prior detailed characterization in the *RHD* literature, RhesusBase, or ISBT allele tables, ensuring that the study focused on functionally novel or incompletely understood regulatory variants.

### sgRNA cloning and plasmid preparation

Single-guide RNAs (sgRNAs) targeting AG-prioritized non-coding variants were designed based on the human reference genome GRCh38/hg38 and are described by variant name throughout the study. Depending on target nucleotide identity and base-editor constraints, sgRNAs were designed to bind either the forward or reverse DNA strand. Potential off-target sites were evaluated using Cas-OFFinder, and guides with no predicted high-risk off-target matches were selected.

sgRNA spacers (20 nt) were cloned into a U6-sgRNA(F+E) empty vector, in which sgRNA expression is driven by a U6 promoter (Addgene: #59986). The backbone vector was digested with BbsI-HF, and complementary oligonucleotides encoding each spacer were annealed and ligated into the linearized vector using T4 DNA ligase under standard conditions. Ligation products were transformed into chemically competent *E. coli* (DH10B), and transformants were selected on ampicillin-containing agar plates. Correct sgRNA insertion and sequence integrity were confirmed by whole-plasmid sequencing prior to downstream experiments.

### CRISPR-nCas9-mediated cytosine base editing and measurement of editing efficiency at the DNA level

To introduce the target SNPs (chr1:25272422 C>T and chr1:25272434 G>A), we employed BE4max (Addgene: #112094), a fourth-generation cytosine base editor consisting of a rat APOBEC1 cytidine deaminase fused to *Streptococcus pyogenes* Cas9 nickase (nCas9, D10A mutation) and two copies of uracil glycosylase inhibitor (UGI). The BE4max system enables efficient C·G to T·A base pair conversion without generating double-strand breaks. The dual UGI domains effectively inhibit cellular uracil DNA glycosylase (UNG), thereby minimizing undesired C to G or C to A byproducts and improving product purity. For CRISPR-nCas9-mediated cytosine base editing, two sgRNAs were designed to introduce the target SNPs using the BE4max system. To generate the G>A substitution at chr1:25272434, an sgRNA targeting the antisense strand (5’-GCCTTGCAGCCTGAGATAAG-3’) was designed, allowing BE4max to convert the complementary cytosine to thymine on the antisense strand. To generate the C>T substitution at chr1:25272422, an sgRNA targeting the sense strand (5’-CAGCCTTGCAGCCTGAGAT-3’) was designed. Oligonucleotides for sgRNA cloning were synthesized as follows: for the first sgRNA, forward oligo 5’-CACCGCTTATCTCAGGCTGCAAGGC-3’ and reverse oligo 5’-AAACGCCTTGCAGCCTGAGATAAGC-3’; for the second sgRNA, forward oligo 5’-CACCGCAGCCTTGCAGCCTGAGATA-3’ and reverse oligo 5’-AAACTATCTCAGGCTGCAATTCTGC-3’. The annealed oligonucleotides were cloned into the sgRNA expression vector.

Base editing efficiency was quantified in K562 cells at the genomic DNA level. Cells were co-transfected with BE4max plasmid (750 ng), sgRNA expression plasmid (250 ng), and a GFP reporter plasmid (2 µg) to enable fluorescence-activated cell sorting (FACS) enrichment of successfully transfected cells. Non-targeting sgRNA-transfected cells were included as negative controls to account for background editing. Two independent transfections were performed for each sgRNA to ensure reproducibility. GFP-positive cells were sorted at 48 h post-transfection, and genomic DNA was extracted at 72 h for downstream analysis. Editing efficiency was assessed by Sanger sequencing followed by deconvolution analysis using EditR or ICE software.

At 72 hours post-transfection, GFP-positive cells were isolated by fluorescence-activated cell sorting. Genomic DNA was extracted from approximately 0.5 × 10□ cells using the QIAamp DNA Micro Kit. Target loci were PCR-amplified (100–200 bp amplicons; 30 cycles) and subjected to next-generation deep sequencing (Plasmidsaurus, Premium PCR). Editing efficiency was calculated as the percentage of sequencing reads containing base conversions at each nucleotide position. Bystander edits within the BE4max activity window were quantified individually.

### Quantification of *RHD* expression by qPCR

Total RNA was extracted from GFP-enriched K562 cells 72 hours post-transfection using TRIzol, followed by column purification with the PureLink RNA Mini Kit and on-column DNase treatment. RNA was isolated from approximately 2 × 10^6^ cells per sample, and 1 µg of total RNA was reverse-transcribed using the High-Capacity cDNA Reverse Transcription Kit.

Quantitative PCR was performed using SYBR Green chemistry (Bio-Rad, Hercules, USA) with primers specific to the full-length *RHD*-201 transcript, spanning exon–exon junctions. GAPDH was used as the internal normalization control. The following primers were used: *RHD* forward 5’-CCCACAGCTCCATCATGGG-3’, *RHD* reverse 5’-GCCAATCATGCCATTGCCGGC-3’, GAPDH forward 5’-GACAGTCAGCCGCATCTTCT-3’, GAPDH reverse 5’-GCGCCCAATACGACCAAATC-3’. Each biological replicate was analyzed in three technical replicates, and relative expression levels were calculated as fold change compared with the non-targeting sgRNA control using the ΔΔCt method.

### Flow cytometric analysis of surface *RHD* expression

Surface *RHD* expression on K562 cells was measured by flow cytometry using indirect immunofluorescence. Non-targeting sgRNA control and SNP-edited K562 cells were harvested during log-phase growth, washed once with PBS containing 2% fetal bovine serum (FBS), and resuspended at 5 × 10□–1 × 10□ cells per 100 µL. Cells were incubated with Seraclone® Anti-D (RH1) 226 human IgM antibody (Bio-Rad, #802042100) for 30 minutes on ice, washed twice, and then incubated for 30 minutes on ice with an IgM-specific, Fc5µ-fragment–specific Alexa Fluor 647–conjugated secondary antibody (Jackson ImmunoResearch, #115-605-020) at a 1:500 dilution. Cells were washed twice and resuspended in PBS + 2% FBS for acquisition. Unstained and secondary-only controls were included for each cell line, and *RHD*-positive red blood cells were stained in parallel as a positive control. Data were acquired on a Cytek Northern Lights (3-laser) spectral flow cytometer with autofluorescence extraction enabled, using a single unstained K562 sample as the spectral reference. Data were analyzed in FlowJo v10.1.00.

## Results

### AG defines predictable regions at the *RHD* locus and establishes a baseline reference for *RHD* expression prediction

We first conducted a systematic prediction and perturbation analysis of the *RHD* locus using AG. Given that the *RHD* gene gives rise to multiple transcripts, we extracted a set of *RHD*-associated transcripts at the transcript level and selected transcript 201 (https://www.ensembl.org/Homo_sapiens/Gene/Summary?db=core;g=ENSG00000187010;r=1:25272393-25330445) as the primary analysis target. This transcript was used as a unified reference for downstream expression readouts and functional interpretation (Figure 1).

**Figure 1.**
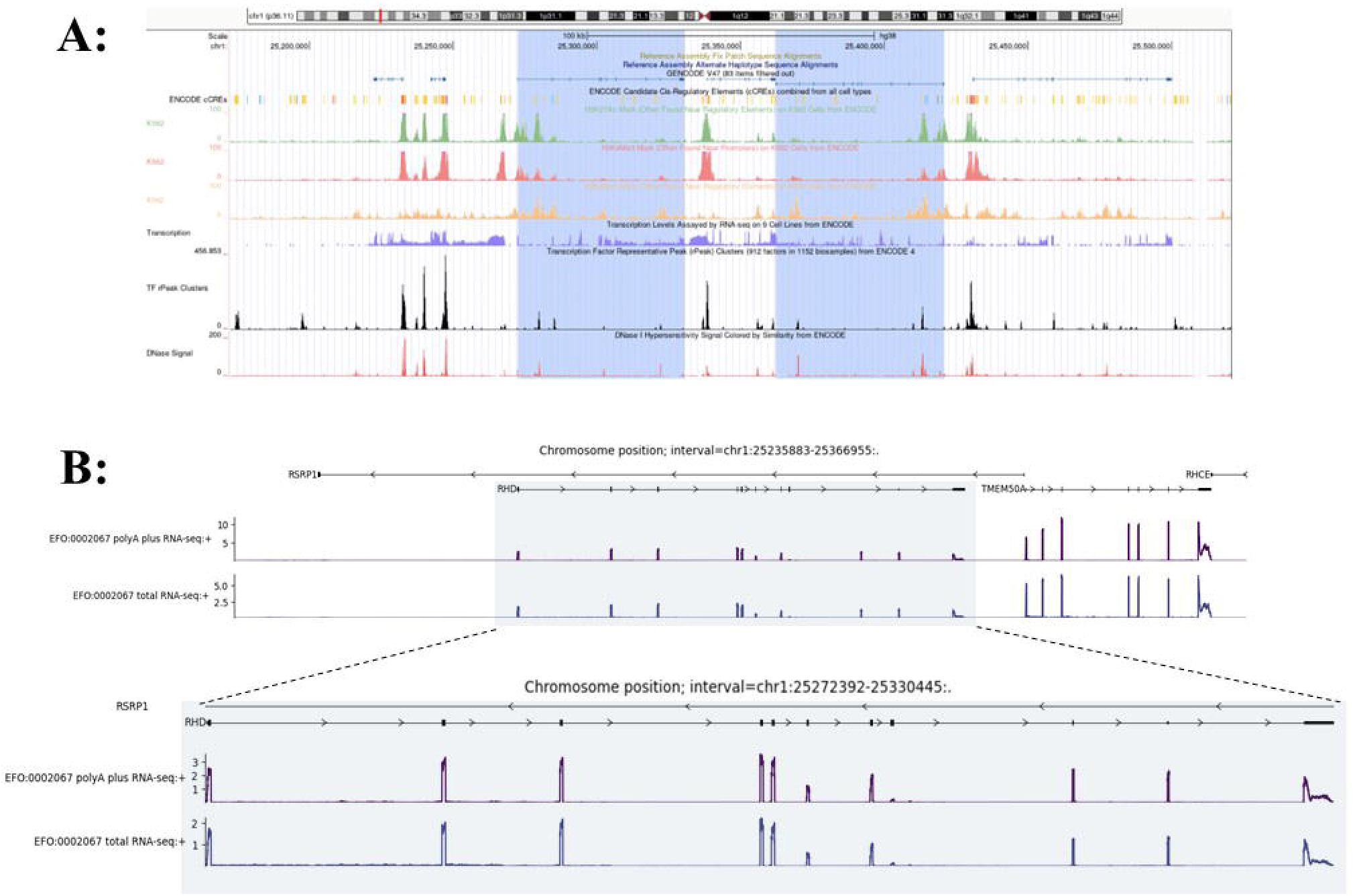
Overview of the *RHD* locus and AG baseline expression prediction. (A) Genomic overview of the *RHD* locus and neighboring genes on chromosome 1 (hg38), showing gene structure together with ENCODE-annotated candidate cis-regulatory elements distributed across upstream, intragenic, and downstream non-coding regions that collectively define the AG screening window. Tracks display K562 cell histone modification ChIP-seq signals for H3K4me3, H3K4me1, and H3K27ac, DNase I hypersensitivity, and ENCODE transcription factor representative peak (rPeak) clusters. These features reveal regions with enriched regulatory activity near the annotated transcription start site and within intragenic intervals at the *RHD* locus. The integrated chromatin accessibility and transcription factor binding patterns informed the selection of candidate intragenic regulatory regions. (B) AG-predicted wild-type RNA expression profile of the *RHD* gene in K562 cells. This prediction represents the baseline and unperturbed expression state used as the reference for subsequent in silico variant perturbation and regulatory effect analyses.

Under the input-length constraints supported by AG (i.e., fixed-size sequence windows only), we defined two major categories of regions for focused screening around the *RHD* locus:

1. a proximal regulatory region spanning ±10 kb around the TSS;
2. intragenic non-coding regions, aimed at assessing the potential contribution of promoter, intronic enhancers, silencers, or splicing-related regulatory elements to *RHD* expression.

For *RHD* expression prediction, RNA-seq was used as the output modality, and the predicted expression derived from the wild-type reference sequence served as the baseline. Under this configuration, the reference prediction score at the *RHD* locus was 0.4054, which was used as the unified control for all subsequent perturbation effects, including deletions and SNPs (Figure 2).

**Figure 2.**
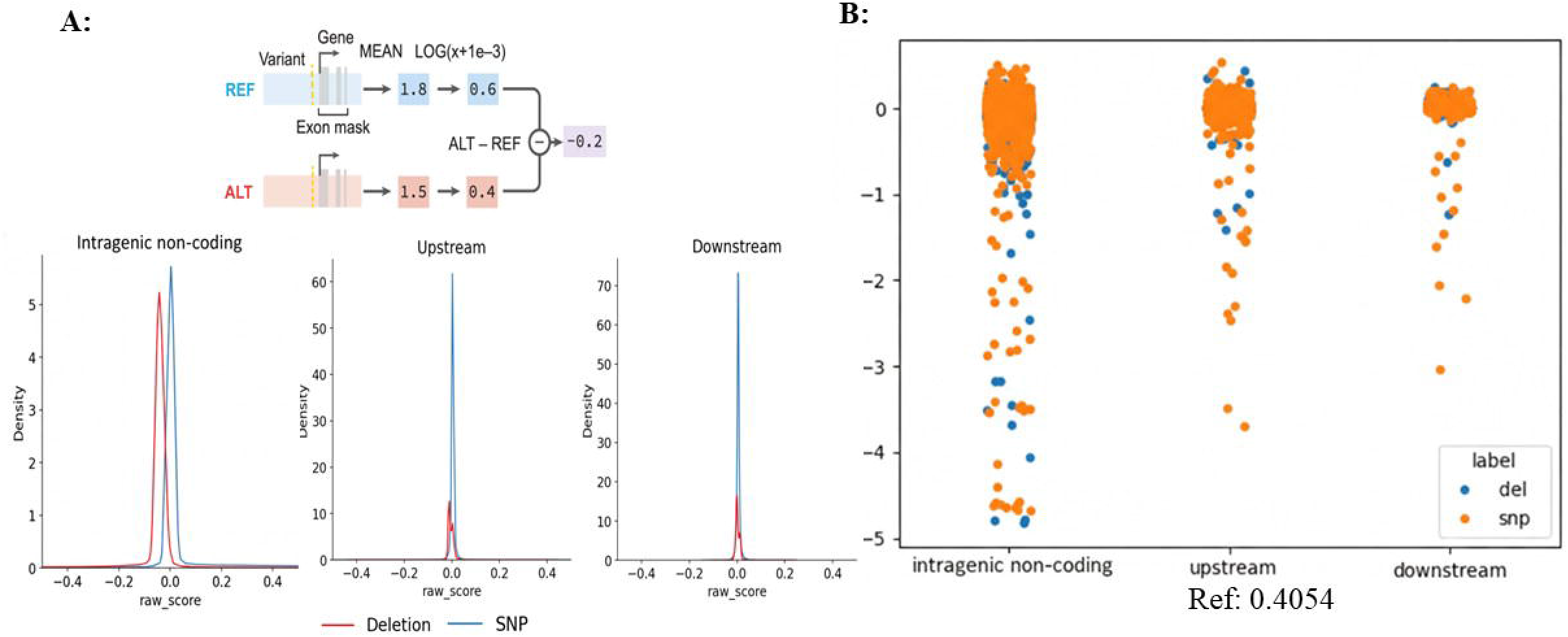
AG RNA-seq–based variant effect scoring scheme and predicted regulatory landscape across the *RHD* locus. (A) Schematic illustration of the AG RNA-seq–based variant effect scoring framework, in which the predicted impact of a variant is quantified as the log-fold change (LFC) in gene expression between the alternative (ALT) and reference (REF) alleles. Positive values indicate predicted upregulation, whereas negative values indicate predicted downregulation of *RHD* expression relative to the reference state. (B) AG-predicted variant effect landscape across the *RHD* locus. Predicted effects of SNPs and deletion perturbations are shown for variants distributed across the *RHD* gene (including intragenic non-coding regions) as well as the 10 kb upstream and downstream flanking regions. The baseline predicted expression level for the *RHD* reference allele (REF) is 0.4054, and all variant scores are reported as LFC values relative to this reference.

### Key regulatory signals are enriched in promoter-proximal and intragenic non-coding regions

Next, we performed two types of *in silico* perturbations across *RHD* intragenic non-coding regions: segmental deletions and single-nucleotide substitutions (SNPs), and quantified RNA-seq–based expression changes using AG. Score directionality was defined relative to the unperturbed state, where values < 0 indicate downregulation of *RHD* expression, and lower scores correspond to stronger repression.

Comparisons showed that perturbations within intragenic non-coding regions were more likely to induce *RHD* downregulation than perturbations within the proximal ±10 kb region, suggesting that key regulatory elements controlling *RHD* expression are more likely to be enriched within introns rather than being restricted to the immediate promoter vicinity.

We further scanned intragenic non-coding regions in a segmented manner using deletion-based perturbation analysis and identified multiple candidate functional segments, among which regions proximal to the promoter exhibited the most pronounced predicted downregulatory effects on *RHD* expression. s. Specifically, among a set of candidate deletion windows spanning promoter-proximal and intragenic non-coding regions (e.g., chr1:25272393–25272547, chr1:25272696–25284572, chr1:25284760–25290640, chr1:25290792–25300945, chr1:25301094–25301519, and chr1:25301687–25303321), AG predictions indicated that several windows exhibited pronounced suppressive effects on *RHD* expression. Overall, deletion-based perturbation analysis across these regions revealed heterogeneous regulatory impacts, highlighting multiple genomic segments with strong predicted downregulatory potential (Figure 3). These regions were therefore prioritized for subsequent SNP-level fine-mapping and experimental validation. SNP-based analyses within promoter-proximal and intragenic non-coding regions yielded a set of candidate variants, which were aggregated by genomic intervals and cross-validated against deletion-based regional signals to increase confidence in identifying regions with consistent predicted regulatory effects.

**Figure 3.**
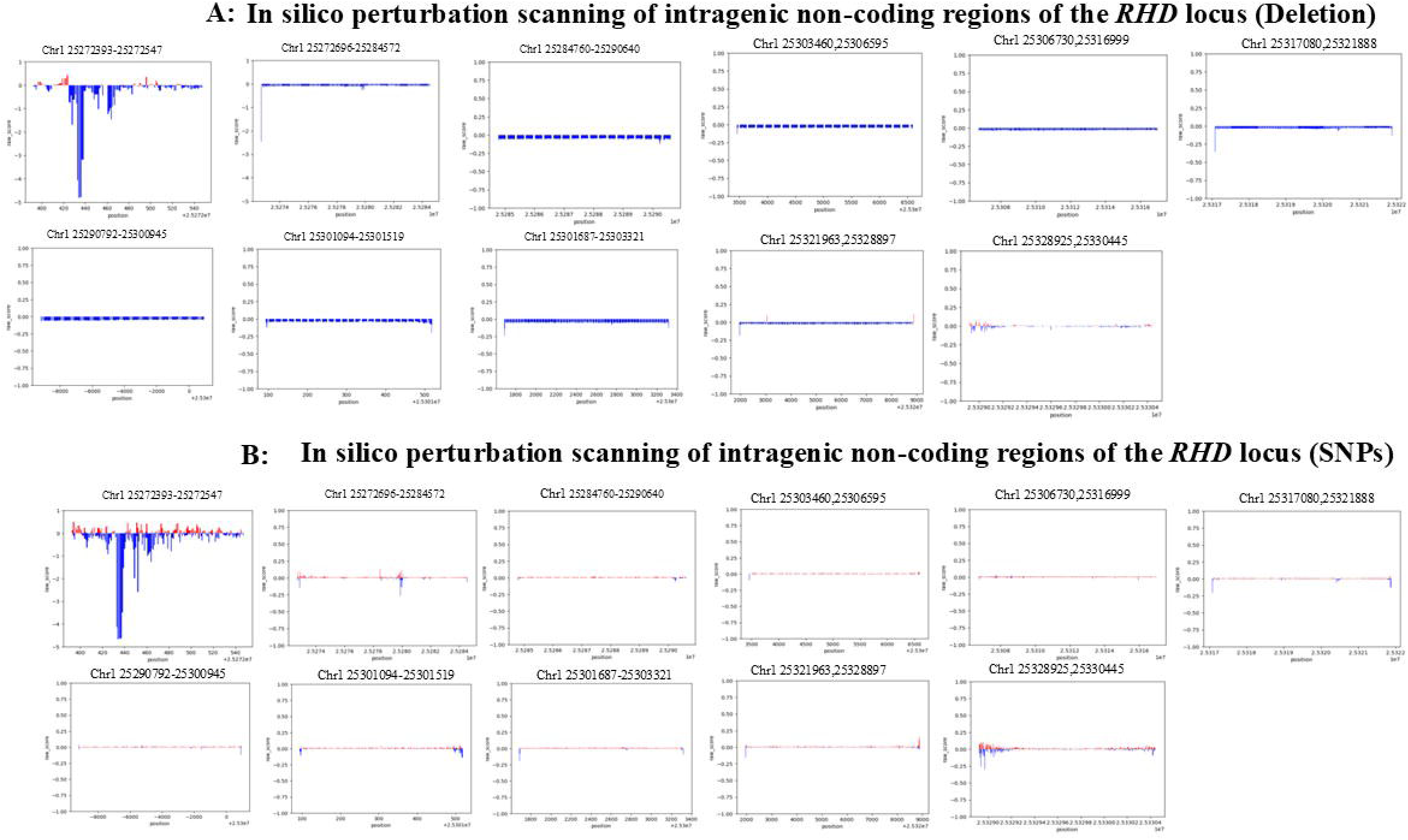
In silico perturbation scanning of *RHD* regulatory regions identifies strong repressive elements within promoter-proximal and intragenic non-coding regions. (A) Deletion-based in silico perturbation scanning across defined promoter-proximal and intragenic non-coding regions at the *RHD* locus. Consecutive short genomic segments were systematically deleted, and the predicted effect of each deletion on *RHD* expression was quantified. The genomic interval chr1:25272393–25272547, which overlaps the annotated *RHD* promoter and immediately upstream promoter-proximal region, shows pronounced negative effects upon deletion, consistent with the presence of essential transcriptional regulatory elements. Additional strong downregulatory signals are observed within intragenic non-coding regions, particularly within the first intron. (B) SNP-based in silico perturbation scanning across the same promoter-proximal and intragenic non-coding regions. Each data point represents the predicted change in *RHD* expression associated with a single-nucleotide substitution at the corresponding genomic position. Consistent with the deletion analysis, single-nucleotide perturbations within the promoter-proximal interval (including chr1:25272393–25272547) and selected intragenic non-coding regions result in substantial predicted downregulation of *RHD* expression. For both panels, the y-axis represents predicted expression changes relative to the unperturbed reference sequence, with values below zero indicating repression of *RHD* expression and lower values corresponding to stronger suppressive effects. Together, these analyses demonstrate that both promoter-proximal regulatory sequences and intragenic non-coding elements—particularly those located within the first intron—play critical roles in maintaining *RHD* transcriptional activity.

To obtain a global view of regulatory sensitivity across the *RHD* locus, we first performed in silico perturbation scanning across an extended genomic region encompassing the *RHD* gene body and its surrounding regulatory regions (Figure 4). This analysis revealed marked heterogeneity in regulatory effects across different genomic segments, with multiple regions exhibiting predicted downregulatory impacts on *RHD* expression, indicating that promoter-proximal and intragenic regions play important roles in the regulation of *RHD* expression. Building on this global analysis, we next focused on the ±10 kb region flanking the TSS of *RHD* to more precisely characterize the contribution of proximal regulatory regions to *RHD* expression (Figure 5). SNP-based perturbation results within this window revealed a clear positional gradient, whereby sequence alterations located closer to the TSS were more likely to induce stronger *RHD* downregulation. Consistent with this observation, a complementary deletion-based evaluation framework implemented for the same region further supported the functional importance of promoter and TSS-proximal regions in regulating *RHD* expression.

**Figure 4.**
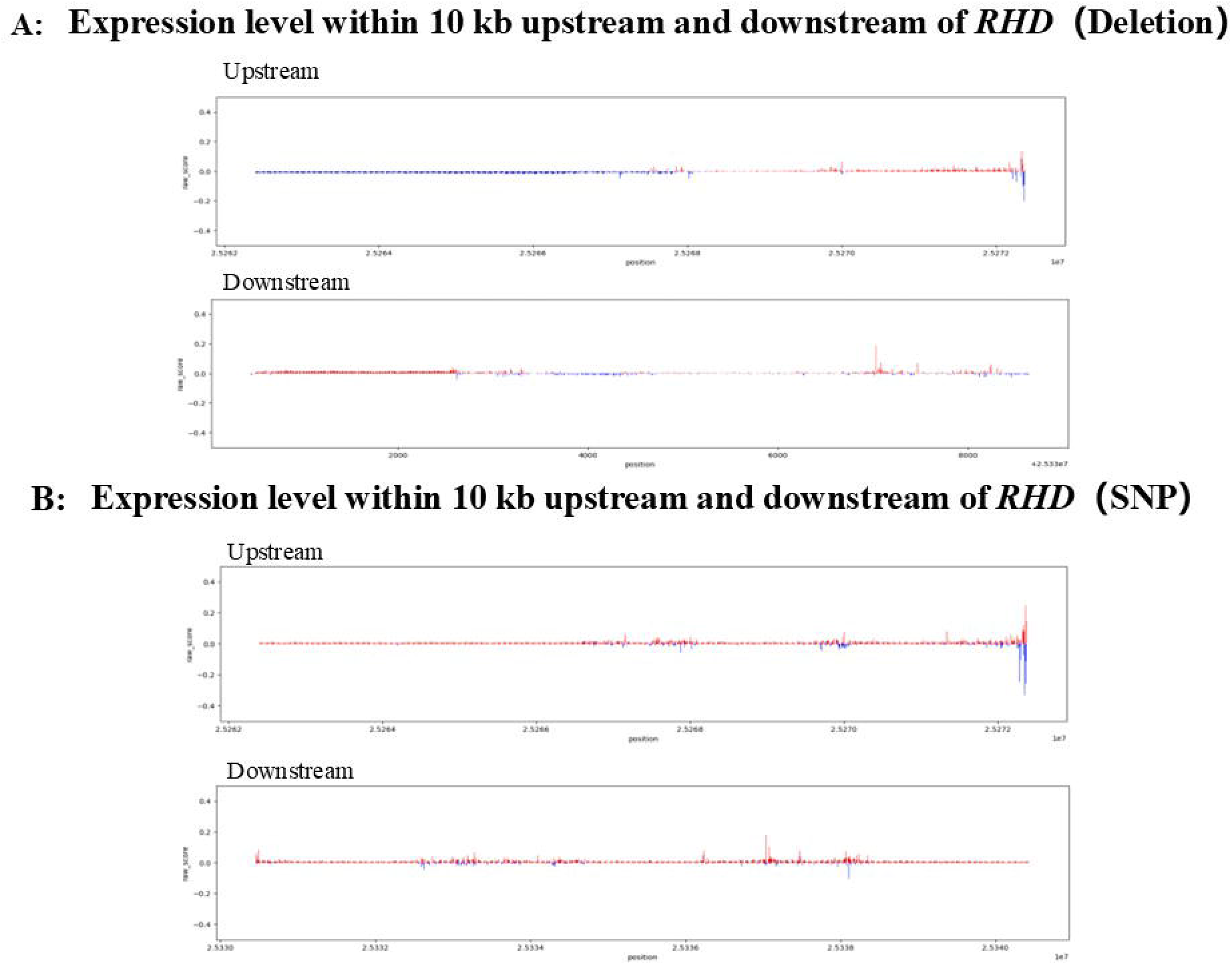
Perturbation analysis of the ±10 kb promoter-proximal region of the *RHD* locus reveals position-dependent downregulatory effects. (A) SNP-based perturbation analysis of candidate variants located within the ±10 kb region flanking the *RHD* promoter. Predicted expression changes are shown as log-fold changes relative to the unperturbed reference state. (B) Deletion-based perturbation analysis across the same ±10 kb promoter-proximal region, providing a complementary assessment of regional regulatory sensitivity. Across both panels, variants located closer to the promoter exhibit more pronounced predicted downregulation of *RHD* expression, indicating that promoter-proximal regulatory regions play a functionally important role in controlling *RHD* expression. Ranked candidate variant sets (e.g., top-ranked variants) are provided to facilitate experimental prioritization.

**Figure 5.**
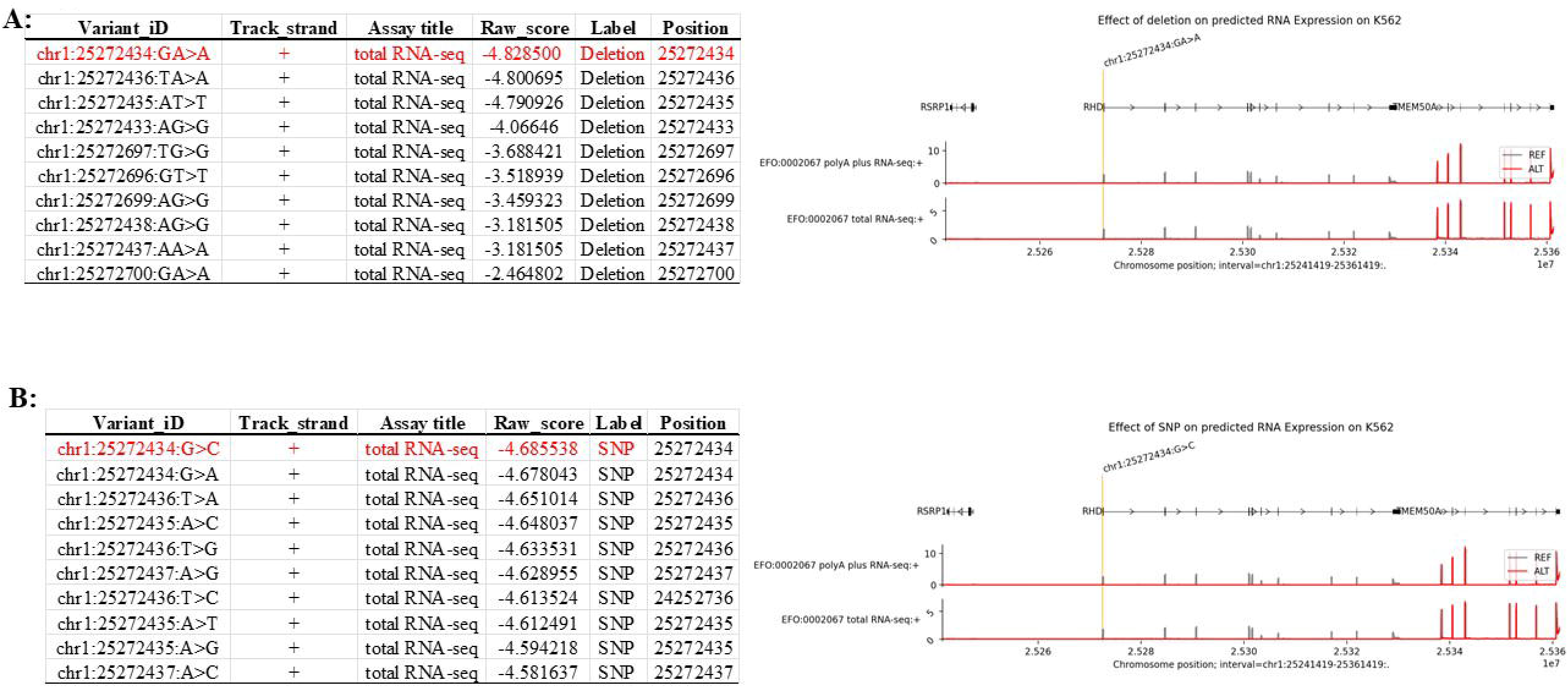
AG-guided prioritization of high-impact regulatory variants at the *RHD* locus. (A) Top-ranked non-coding regulatory variants identified by AG based on predicted suppressive effects on *RHD* expression. Candidate sites represent the highest-scoring variants derived from deletion- and SNP-based perturbation analyses and are displayed within their local genomic sequence contexts. The top 10 variants from each perturbation mode were selected to define a high-confidence candidate set for downstream investigation. (B) (Top) Predicted impact of the highest-scoring genomic deletion identified in the deletion-based perturbation analysis at chr1:25272434, representing the strongest predicted downregulatory effect on *RHD* RNA expression among all tested deletions. (Bottom) Predicted impact of the highest-scoring single-nucleotide substitution identified in the SNP-based perturbation analysis at the same genomic position (chr1:25272434), illustrating the strongest predicted effect among all evaluated single-nucleotide variants. These examples are shown to illustrate how AG ranks and models the most influential regulatory perturbations across different variant classes.

In this analysis, we additionally generated ranked candidate sets (e.g., top-10 deletions and top-10 SNPs) to guide experimental prioritization, enabling focused validation of the most likely functional sites within a manageable experimental scale.

### Functional validation of AG-prioritized non-coding SNPs through base-editing technology

Based on the in silico scanning results described above, we selected high-priority non-coding SNPs located within promoter-proximal and intragenic non-coding regions of *RHD* for experimental validation. Figure 6 explicitly shows two candidate editing sites near the transcription initiation region—chr1:25272422 (c.1-83 C>T) and chr1:25272434 (G>A)—together with their surrounding sequence contexts, which were used to guide sgRNA design and editing-window coverage assessment. GNOMAD was used to assess allele frequencies; the variants are present at very low frequency (allele frequency < 1 × 10□□), consistent with potential phenotypic impact, but they have not been previously characterized or reported in the literature. To quantitatively assess *RHD* phenotypes at the transcriptional level, we employed a qPCR primer system targeting the full-length *RHD*-201 transcript, allowing *RHD* expression changes before and after base editing to be compared under a unified transcript definition.^26,27^ This strategy avoids interpretational bias arising from changes in transcript isoform proportions and enables robust evaluation of regulatory effects induced by non-coding variants (Figure 7). Both variants, chr1:25272434 (G>A) and chr1:25272422 (c.1-83 C>T), resulted in a marked reduction in *RHD* gene expression, providing experimental validation of the predictive performance of the AG model. To further assess whether transcriptional changes induced by non-coding variant editing translate into alterations at the protein level, we examined surface *RHD* expression by flow cytometry (Figure 8). K562 cells edited at chr1:25272422 (C>T) and chr1:25272434 (G>A), together with non-targeting sgRNA controls, were stained with a human anti-D IgM antibody and analyzed by spectral flow cytometry. Consistent with the qPCR results, cells harboring the chr1:25272434 (G>A) edit exhibited a clear leftward shift in fluorescence intensity compared with control cells, indicating reduced surface *RHD* expression. In contrast, editing at chr1:25272422 resulted in a more modest decrease in *RHD* signal, consistent with its weaker transcriptional effect. These results demonstrate that AG-prioritized non-coding variants not only modulate *RHD* transcript abundance but also produce corresponding changes at the cell-surface antigen level, providing functional validation at both transcriptional and phenotypic layers.

**Figure 6.**
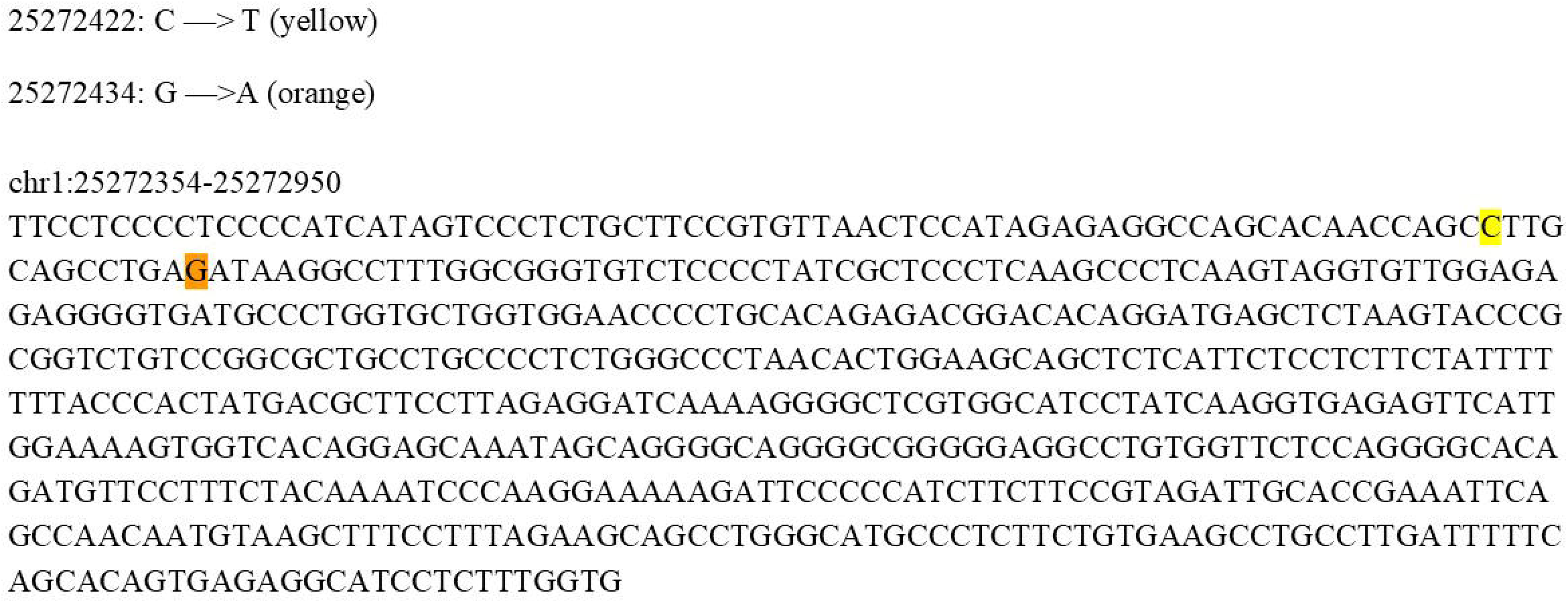
Genomic locations of candidate non-coding variants selected for base-editing validation at the *RHD* locus. Sequence view of the genomic region surrounding the *RHD* transcription start site (chr1:25272354–25272950, hg38), highlighting two AG-prioritized non-coding variants selected for experimental validation: chr1:25272422 (c.1–83 C>T, highlighted in yellow) and chr1:25272434 (G>A, highlighted in orange). The displayed nucleotide sequence provides the exact genomic context used for sgRNA design and evaluation of the BE4max editing window, including the relative positions of target bases and potential bystander nucleotides. Both variants reside within a promoter-proximal regulatory region predicted by AG to exert suppressive effects on *RHD* expression.

**Figure 7.**
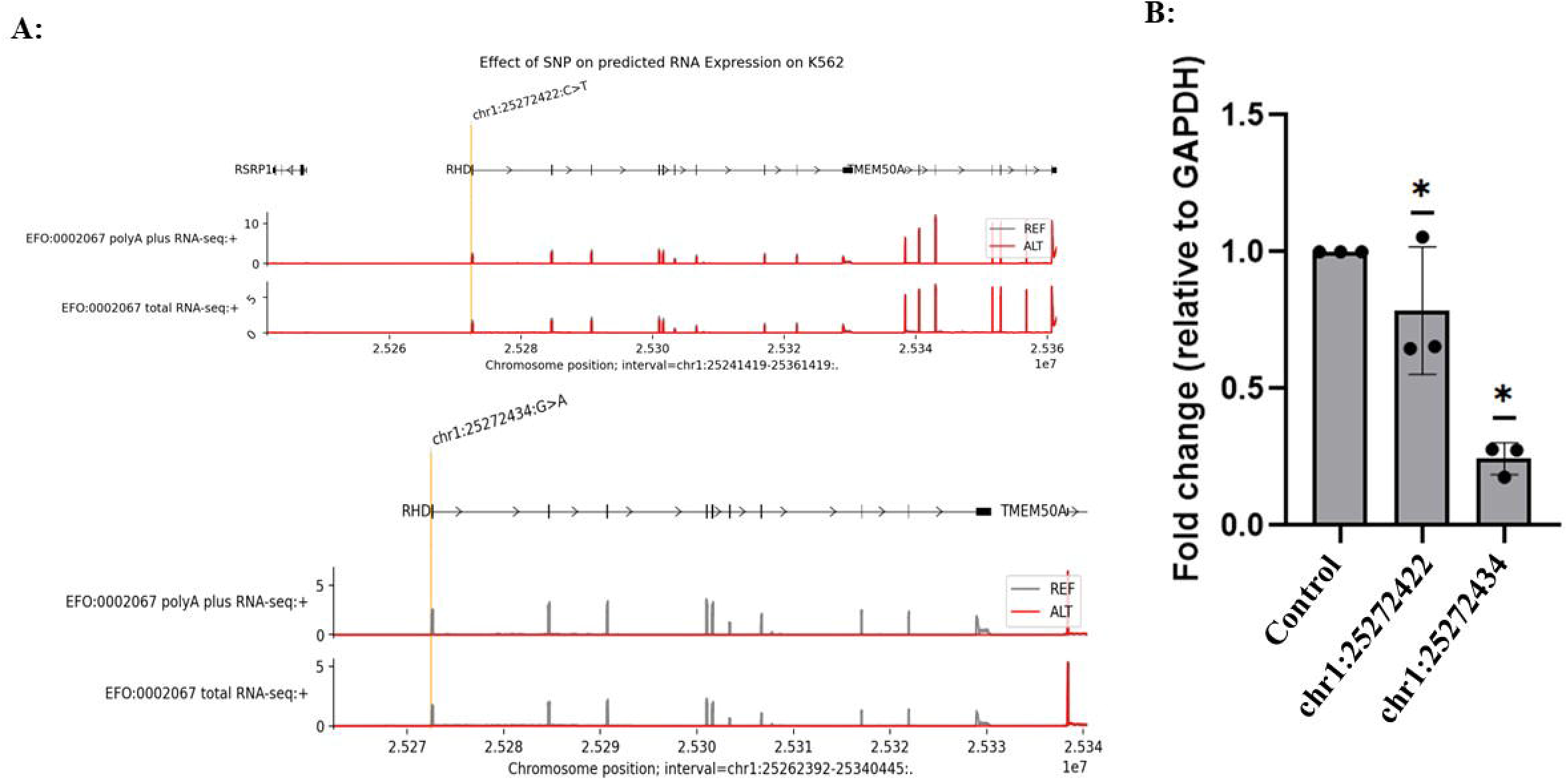
Effects of CRISPR-mediated base editing at selected non-coding variants on *RHD* transcript expression. **(A)** AG-predicted RNA-seq coverage profiles across the *RHD* locus for the reference allele (REF, gray) and alternative alleles (ALT, red). A zoomed-in view highlights distinct predicted regulatory effects of two promoter-proximal variants: the C>T substitution at chr1:25272422 is associated with a modest reduction in predicted *RHD* expression, whereas the G>A substitution at chr1:25272434 produces a pronounced downregulatory effect, consistent with its higher AG score. **(B)** Experimental validation of *RHD* expression changes following base editing in K562 cells, measured by qPCR targeting the full-length *RHD*-201 transcript. Relative expression levels are shown for cells edited at chr1:25272422 and chr1:25272434, normalized to non-targeting sgRNA controls. Editing at chr1:25272434 results in a significant reduction of *RHD* transcript abundance, whereas editing at chr1:25272422 leads to a comparatively weaker effect, mirroring the differential regulatory impact predicted by AG.

**Figure 8.**
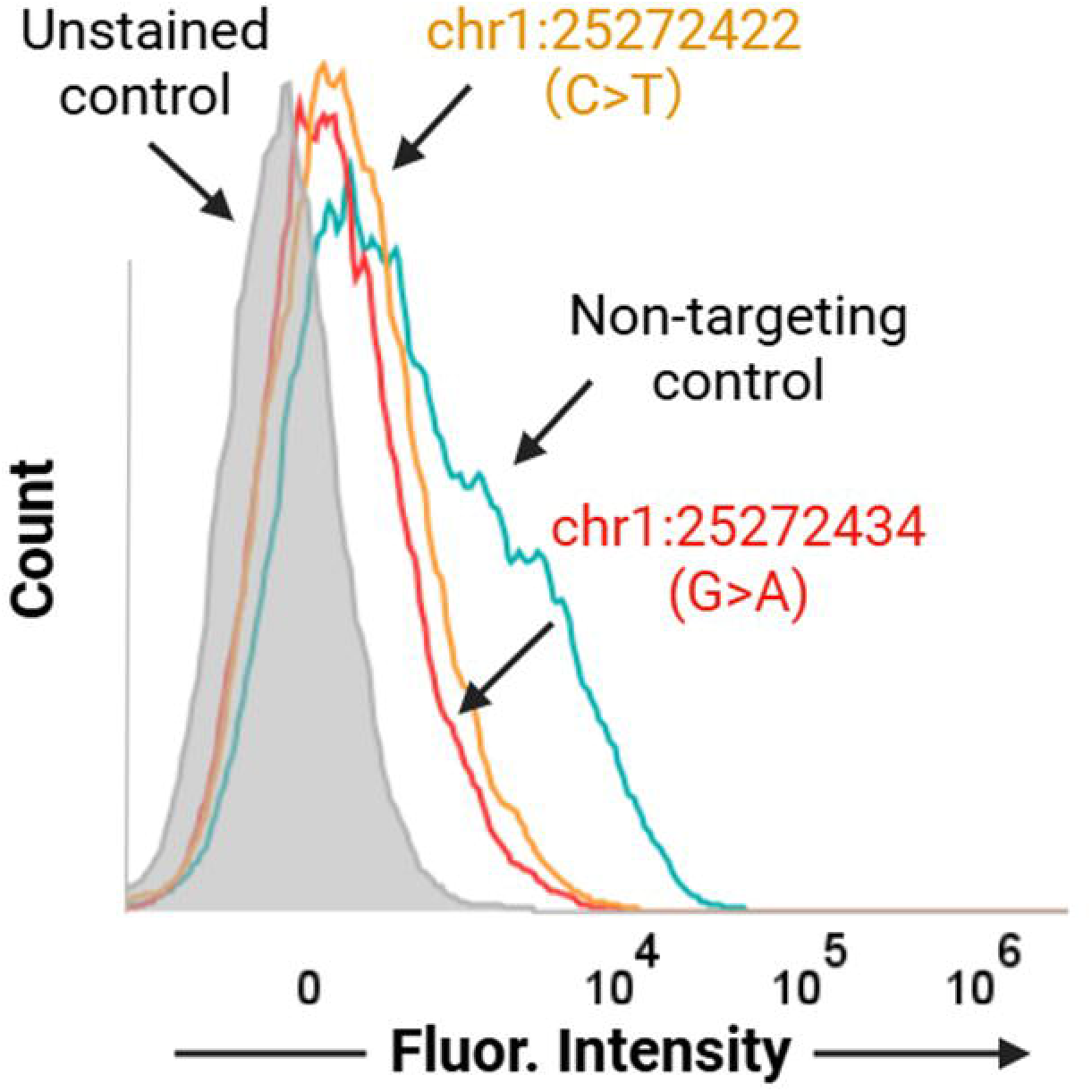
Flow cytometric detection of surface *RHD* expression in K562 cells following SNP editing. Representative overlaid histograms showing surface *RHD* staining on K562 cells measured by spectral flow cytometry using a human anti-D IgM primary antibody and an IgM-specific Alexa Fluor 647 secondary antibody. Unstained cells (gray) define baseline fluorescence. Non-targeting sgRNA control cells (teal) show a clear rightward shift, indicating detectable surface *RHD* expression. In contrast, cells edited at *RHD*-associated noncoding variants chr1:25272422 (C>T, orange) and chr1:25272434 (G>A, red) exhibit reduced fluorescence intensity relative to the non-targeting control, consistent with decreased surface *RHD* expression. Data are gated on singlet cells; histograms are normalized to mode and displayed on a biexponential scale.

## Discussion

In this study, we combined AI-based regulatory variant prediction with targeted base editing to systematically interrogate non-coding determinants of *RHD* expression. Our results provide experimental evidence that deep-learning–guided prioritization can accurately identify functionally relevant regulatory variants and substantially narrow the search space traditionally addressed by large-scale CRISPR screening or population-level association studies.

Notably, the concordance observed between AG predictions and wet-lab outcomes supports the biological validity of this integrative framework and highlights its potential utility for dissecting regulatory mechanisms in clinically significant blood group genes. AG-predicted regulatory hotspots across the *RHD* locus also showed strong concordance with regions previously highlighted by manual epigenetic annotation, particularly within the promoter and its proximal regions as well as intragenic non-coding regions. Across both analytical frameworks, variants located in these regions consistently emerged as key modulators of *RHD* expression, underscoring the importance of non-coding regulatory architecture in controlling RHD transcription.

A central finding of this work is that non-coding variants within promoter-proximal and intragenic non-coding regions exert disproportionately strong effects on gene expression. Such SNP-mediated regulatory perturbations are increasingly recognized as key contributors to disease susceptibility, therapeutic response, and mechanistic heterogeneity across diverse biological contexts.^28-31^ AG-based in silico perturbation analyses consistently predicted marked downregulation of *RHD* expression following deletions or SNPs in these regions, whereas variants located further upstream or downstream showed comparatively modest effects. These predictions were subsequently confirmed experimentally: base editing of a high-impact predicted SNP (chr1:25272434 G>A) resulted in efficient editing and measurable transcriptional consequences, while editing attempts at lower-impact predicted sites failed to produce meaningful changes. Together, these results demonstrate the ability of AI-based models to prioritize functionally relevant regulatory variants that are difficult to infer from sequence context alone.

The functional importance of intragenic non-coding regulatory regions observed in this study adds to a growing body of evidence indicating that non-coding regions within gene bodies can play active roles in fine-tuning transcriptional output. Rather than serving solely as passive intervening sequences, such regions may contribute to transcriptional regulation through diverse mechanisms. Our findings further suggest that variation within intragenic non-coding regions represents an underappreciated source of regulatory diversity, which may help explain why individuals with identical coding sequences can nonetheless exhibit divergent *RHD* serologic phenotypes, a long-standing observation in transfusion medicine.^15,32,33^

The base editing experiments further revealed an important and often underappreciated aspect of regulatory variant interrogation: unintended or bystander edits can themselves be informative rather than purely confounding. In our study, editing at chr1:25272434 G>A was accompanied by nearby base conversions, each of which was independently predicted by AG to reduce *RHD* expression. Rather than obscuring interpretation, this clustering of functionally consistent edits reinforces the notion that regulatory regions often act as integrated modules, where multiple adjacent nucleotides collectively contribute to transcriptional control. From a mechanistic perspective, such regions may correspond to transcription factor binding motifs, chromatin accessibility hotspots, or RNA regulatory elements that are sensitive to subtle sequence perturbations.^34,35^

These observations have broader implications for experimental design. Traditional CRISPR knockout approaches typically disrupt large genomic segments, making it difficult to disentangle the contributions of individual nucleotides.^36^ In contrast, base editing allows precise, single-nucleotide–level perturbation while preserving local chromatin architecture.^37^ When guided by AI-based predictions, this precision enables efficient causal testing of candidate regulatory variants, transforming what would otherwise require extensive screening into a focused validation strategy. This approach is particularly advantageous for genes such as *RHD*, where regulatory complexity and genomic homology present significant technical challenges. Our results also highlight the limitations of relying exclusively on coding-region analyses for molecular blood group typing. Although coding variants in *RHD* have been extensively cataloged and successfully incorporated into clinical genotyping algorithms, they do not fully capture the spectrum of *RHD* expression variability observed in practice. The identification of functional non-coding variants provides a complementary layer of information that may help resolve discordant genotyping–phenotyping cases, particularly in individuals with weak D or DEL phenotypes.^38,39^ In this context, AI-guided regulatory variant analysis has the potential to enhance the precision of genomics-based transfusion medicine.

Despite these strengths, several limitations of the present study warrant consideration. First, our experimental validation was performed in erythroid cell line models, which, while highly informative, cannot fully recapitulate the complexity of in vivo erythropoiesis. Regulatory effects may vary across differentiation stages or in primary erythroid progenitors, and future studies incorporating primary cells or in vivo models will be essential to extend these findings. Second, while AG demonstrated strong predictive performance for *RHD*, AI models are inherently dependent on the quality and diversity of their training data. Regulatory elements that are highly cell-type–specific or poorly represented in existing functional genomics datasets may be more difficult to predict accurately.

Additionally, the scope of base editing validation in this study was necessarily limited to a subset of prioritized variants. Although this reflects a deliberate strategy to demonstrate feasibility and predictive accuracy, comprehensive mapping of all potential regulatory variants within the *RHD* locus will require further investigation. Integration with complementary approaches, such as chromatin accessibility profiling or allele-specific expression analyses, may provide additional mechanistic insight and help refine predictive models.

Looking forward, the framework established here offers several avenues for expansion. Applying this AI-guided strategy to larger cohorts and additional blood group genes could systematically uncover regulatory variants that contribute to transfusion-related risk. Moreover, coupling regulatory variant prediction with emerging long-read sequencing technologies may improve variant detection in highly homologous genomic regions such as the *RH* locus. Beyond transfusion medicine, this approach is broadly applicable to other clinically relevant genes where non-coding variation plays a critical but underexplored role.

In conclusion, our study demonstrates that AI-guided prioritization of non-coding regulatory variants, combined with precise base editing validation, provides a powerful and scalable alternative to traditional large-scale screening approaches. By revealing previously unrecognized regulatory determinants of *RHD* expression, this work advances mechanistic understanding of *RHD* variability and establishes a generalizable framework for decoding functional non-coding variation in human disease and clinical genetics.

## ACKNOWLEDGEMENTS

This work was supported by the National Institutes of Health (NIH), grant number **P01HL158688**.

